# Inhibition of Cyp1a Protects Mice against Anthracycline Cardiomyopathy

**DOI:** 10.1101/2024.04.10.588915

**Authors:** Jing Liu, Casie Curtin, Rahul Lall, Sarah Lane, Jakob Wieke, Abul Ariza, Leinal Sejour, Ioannis Vlachos, Beshay N. Zordoky, Randall T Peterson, Aarti Asnani

## Abstract

**Background:** Anthracyclines such as doxorubicin (Dox) are highly effective anti-tumor agents, but their use is limited by dose-dependent cardiomyopathy and heart failure. Our laboratory previously reported that induction of cytochrome P450 family 1 (Cyp1) enzymes contributes to acute Dox cardiotoxicity in zebrafish and in mice, and that potent Cyp1 inhibitors prevent cardiotoxicity. However, the role of Cyp1 enzymes in chronic Dox cardiomyopathy, as well as the mechanisms underlying cardioprotection associated with Cyp1 inhibition, have not been fully elucidated.

**Methods:** The Cyp1 pathway was evaluated using a small molecule Cyp1 inhibitor in wild-type (WT) mice, or Cyp1-null mice (*Cyp1a1/1a2^-/-^*, *Cyp1b1^-/-^*, and *Cyp1a1/1a2/1b1^-/-^*). Low-dose Dox was administered by serial intraperitoneal or intravenous injections, respectively. Expression of *Cyp1* isoforms was measured by RT-qPCR, and myocardial tissue was isolated from the left ventricle for RNA sequencing. Cardiac function was evaluated by transthoracic echocardiography.

**Results:** In WT mice, Dox treatment was associated with a decrease in *Cyp1a2* and increase in *Cyp1b1* expression in the heart and in the liver. Co-treatment of WT mice with Dox and the novel Cyp1 inhibitor YW-130 protected against cardiac dysfunction compared to Dox treatment alone. *Cyp1a1/1a2^-/-^* and *Cyp1a1/1a2/1b1^-/-^* mice were protected from Dox cardiomyopathy compared to WT mice. Male, but not female, *Cyp1b1^-/-^* mice had increased cardiac dysfunction following Dox treatment compared to WT mice. RNA sequencing of myocardial tissue showed upregulation of *Fundc1* and downregulation of *Ccl21c* in *Cyp1a1/1a2^-/-^* mice treated with Dox, implicating changes in mitophagy and chemokine-mediated inflammation as possible mechanisms of Cyp1a-mediated cardioprotection.

**Conclusions:** Taken together, this study highlights the potential therapeutic value of Cyp1a inhibition in mitigating anthracycline cardiomyopathy.

## Clinical Perspective

### What is new?

- In this study, we define the role of Cyp1 enzymes in chronic doxorubicin-induced cardiomyopathy using both chemical and genetic approaches in mice.
- Potential mechanisms of Cyp1-mediated cardioprotection include regulation of mitophagy and chemokine-mediated inflammation.
- This study highlights Cyp1a inhibition as a promising therapeutic strategy to mitigate anthracycline cardiomyopathy.

### What are the clinical implications?

Anthracyclines such as doxorubicin are highly effective and often first-line chemotherapies for various malignancies, but up to 20% of patients treated with anthracyclines develop cardiac dysfunction. Several molecular pathways have been implicated in anthracycline cardiomyopathy, but treatments targeting these mechanisms have not been integrated into clinical care. Here we present data supporting inhibition of Cyp1a enzymes as a new therapeutic strategy for doxorubicin cardiomyopathy.

## INTRODUCTION

Cancer outcomes have improved steadily over the past few decades due to earlier detection of malignancy and advances in the development of novel targeted drugs and immunotherapies, contributing to a longer lifespan in cancer survivors.^1^ By the year 2040, the estimated number of cancer survivors is expected to reach 26 million in the United States. Anthracyclines such as doxorubicin (Dox) are well-established antitumor agents and remain among the most effective chemotherapeutic agents for leukemia, lymphomas, sarcomas, and certain types of breast cancer. However, anthracyclines can cause cumulative and dose-dependent cardiomyopathy and heart failure, leading to an increased risk of mortality among cancer survivors and limiting widespread use of these agents.^2^

Anthracycline-associated cardiotoxicity is characterized by left ventricular dysfunction that leads to heart failure in approximately 4% of all patients treated with these agents.^3^ The precise mechanisms underlying anthracycline-associated cardiotoxicity are complex and multifactorial. Proposed molecular pathways include oxidative stress,^4^ mitochondrial iron overload,^5^ topoisomerase II-mediated DNA double-strand breaks,^6^ iron-dependent programmed cell death (ferroptosis),^7^ autophagy,^8^ and disruption of nuclear factor erythroid 2-related factor 2 (NRF2)-,^9^ AMP-activated protein kinase (AMPK)-,^10^ and phosphatidylinositol 3-kinase/protein kinase B (PI3K/Akt)-mediated signaling^11^. However, none of these pathways have been effectively targeted in patients. As an antioxidant and iron chelator, dexrazoxane is the only FDA-approved drug used clinically to prevent Dox cardiomyopathy. It is prescribed to patients at a concentration of 10:1 (dexrazoxane:Dox), underscoring the need for more potent and specific cardioprotectants. Furthermore, the use of dexrazoxane is limited by concerns that it may interfere with Dox’s antitumor effect^12^ and induce secondary malignancies.^13^ Although dexrazoxane’s effects on cancer outcomes remain controversial, the FDA has restricted its use to patients with metastatic breast cancer planned for treatment with at least 300 mg/m^2^ of Dox. Standard heart failure medications such as beta-blockers and renin-angiotensin inhibitors have also been proposed in patients receiving cardiotoxic chemotherapy. Although these appear to be effective once cardiomyopathy is established, randomized controlled trials have not shown a consistent benefit in preventing Dox cardiomyopathy and heart failure.^14–16^ Statins may provide benefit in high-risk groups, but data on their use for primary prevention of Dox cardiomyopathy are conflicting.^17,18^ Thus, there is a need for new mechanism-based cardioprotective strategies to mitigate Dox cardiomyopathy in patients.

Cytochrome P450 (CYP) family 1 enzymes play a critical role in the metabolism of various endogenous and exogenous compounds, including drugs, toxins, and environmental pollutants.^19,20^ The CYP1 subfamily consists of three main isoforms: CYP1A1, CYP1A2, and CYP1B1 in humans and Cyp1a1, Cyp1a2, and Cyp1b1 in mice.^21^ These three enzymes are largely conserved between mice and humans. CYP1A2 is located approximately 25 kb from CYP1A1 on chromosome 15, whereas CYP1B1 is located on chromosome 2.^22^ CYP1A2 is constitutively expressed at high level in human liver, with inducible expression reported in extrahepatic tissues such as pancreas, gastrointestinal tract, brain and heart.^23,24^ In contrast, CYP1A1 and CYP1B1 are primarily expressed by extrahepatic tissues, including the lungs, heart, gastrointestinal tract, kidney and uterus.^25–30^ Notably, CYP1B1 is the most abundantly expressed CYP gene in human hearts.^31^ Cyp1b1 has been reported to contribute to the development of hypertension and renal dysfunction in male mice.^32^ The transcription of CYP1 enzymes is primarily regulated by the aryl hydrocarbon receptor (AhR), a ligand-activated transcription factor that is crucial for metabolizing various foreign substances.^33^ Importantly, Dox has been previously described as a ligand for the AhR.^34^ Following activation, AhR relocates to the nucleus where it forms a complex with the AhR nuclear translocator (ARNT). The AhR/ARNT complex binds to xenobiotic response elements (XRE), inducing expression of *Cyp1a1*, *Cyp1a2*, and *Cyp1b1*. Although a prior study described that activation of the AhR by Dox mediates cytoprotective effects in the heart,^34^ we showed that inhibition of downstream Cyp1 enzymes is protective against Dox cardiac toxicity.^35,36^ These discrepant results could be related to the pleiotropic nature of the AhR, which likely regulates expression of both protective and deleterious pathways other than CYP1 enzymes. We have therefore focused on inhibition of CYP1A enzymes as a more specific target for cardioprotection in the setting of Dox.

In prior work, we performed a phenotypic chemical screen in zebrafish and found that the Cyp1 inhibitor visnagin prevented Dox-induced heart failure.^37^ Through structure-activity relationship studies in zebrafish, we identified the visnagin analog YW-130 (Compound 23 in our prior publication) as preventing acute Dox cardiac toxicity in zebrafish and in mice with high potency.^35^ Moreover, mutation of the *cyp1a* active site in zebrafish protected against Dox-induced heart failure.^36^

In this study, we assessed the role of Cyp1 inhibition in mouse models of chronic Dox cardiomyopathy, which more closely recapitulate the cardiac dysfunction that occurs in patients treated with anthracyclines.^38^ We investigated the functional consequences of inhibiting Cyp1 enzymes using either a small molecule Cyp1 inhibitor in wild-type (WT) mice, or Cyp1-null mice (*Cyp1a1/1a2^-/-^*, *Cyp1b1^-/-^*; and *Cyp1a1/1a2/1b1^-/-^*). Cyp1 inhibition using YW-130, as well as genetic knockout of *Cyp1a* enzymes (*Cyp1a1/1a2^-/-^* and *Cyp1a1/1a2/1b1^-/-^*), protected mice from Dox-induced cardiomyopathy. Our study highlights the potential therapeutic value of Cyp1a inhibition in mitigating anthracycline cardiomyopathy and heart failure.

## METHODS

### Mice

Male and female WT mice on a C57BL/6N background aged 8-10 weeks were purchased from Charles River. *Cyp1a1/1a2^-/-^* and *Cyp1b1^-/-^* mouse founders were generously provided by Dr. Daniel W. Nebert, MD (University of Cincinnati College of Medicine) and Dr. Frank J. Gonzalez, Ph.D. (National Cancer Institute), respectively. To generate *Cyp1a1/1a2/1b1* triple-knockout mice, *Cyp1a1/1a2^-/-^* mice were bred with *Cyp1b1^-/-^* mice for several generations, with genotyping performed at each generation to confirm the strain. All mice were housed in a specific pathogen–free barrier facility at Beth Israel Deaconess Medical Center (BIDMC), maintained in a controlled environment with a 12-hour light/12-hour dark cycle, and provided free access to food and water. Mice displaying signs indicative of a moribund state were euthanized immediately. All animal procedures adhered to the Guide for the Care and Use of Laboratory Animals published by the U.S. National Institutes of Health and were approved by the BIDMC Animal Care and Use Committee.

### Drug preparation and administration

Dox (cat#25316-40-9, Tocris Bioscience) was dissolved in saline at a concentration of 1 mg/ml. YW-130 was initially dissolved in 10% DMSO and then diluted with 90% olive oil to achieve a concentration of 0.25 mg/ml. Small molecule solutions were aliquoted and stored in a −80°C freezer before use, with each aliquot intended for single-use administration. Dox and YW-130 co-treatment was administered via contralateral intraperitoneal (i.p.) injection. Mice aged 12-14 weeks were treated with saline, YW-130, Dox, and Dox with YW-130 every 2 days for 2 weeks, totaling 8 injections.^39^ In knockout models, mice aged 12-14 weeks were treated with saline or Dox once weekly for 4 weeks via intravenous (i.v.) injection in the tail vein, totaling 4 injections.^40^ Echocardiography was performed 7-10 weeks from Dox initiation. Plasma and tissues, such as liver and heart, were collected for further analysis.

### YW-130 pharmacokinetic study

The pharmacokinetics (PK) study was performed by Charles River Laboratories. In brief, YW-130 solution was prepared as described above at a concentration of 0.2 mg/ml. YW-130 1 mg/kg was administered to C57BL/6 mice aged 8-12 weeks by i.p. injection (n=3). Blood samples (∼30 ul) were collected via tail vein at multiple time points: 0.5, 1, 2, 4, 6, 8 and 24 hours post-administration. Concentrations of YW-130 in the plasma were measured using liquid chromatography-mass spectrometry.

### Mouse echocardiography

Conscious mouse echocardiography was performed as previously described.^41^ In brief, the mouse was positioned and fixed on a pre-heated platform to minimize movement. Its limbs were carefully secured, and oxygen was supplied to maintain physiological conditions. Hair on the chest was removed using a hair removal cream (Nair). Conductive gel was applied, and a MS550D ultrasound probe was positioned on the chest area perpendicular to the long axis of the heart. 2D echocardiographic images were then acquired in the parasternal short-axis view. M-mode images were recorded at the level of the papillary muscles to assess cardiac function parameters. Following image acquisition, data analysis was conducted using Vevo 2100 software to quantify left ventricular diameters and fractional shortening (FS%).

### Bulk RNA sequencing

Mice were euthanized one day after the last Dox injection, and heart tissues were collected following perfusion with 20 ml of cold PBS. The total RNA was isolated from heart tissues using Trizol. The quality of the total RNA was assessed using Nanodrop spectrophotometry, and the RNA samples were subjected to library preparation, during which they were converted into cDNA fragments. Subsequently, the cDNA libraries underwent high-throughput sequencing using Illumina. The raw sequencing data were processed to eliminate low-quality reads and adapters, retaining only the clean reads, which were then aligned to the mouse genome using bioinformatics tools. Differential gene expression analysis was conducted to identify genes that were either upregulated or downregulated under different experimental conditions. Functional enrichment analysis including pathway enrichment analysis and gene ontology (GO) enrichment analysis was performed to identify enriched pathways, biological processes, molecular functions, and cellular components among differentially expressed genes.

### RT-qPCR

The total RNA from liver or heart tissues was isolated and qualified as described above (bulk RNA sequencing methods). Total RNA was converted into cDNA fragments using the BioRad iScript Advanced cDNA Synthesis Kit (cat#170-8843). The cDNA fragments were then subjected to qPCR reactions using the PowerUp™ SYBR™ Green Master Mix (cat#A25778) and primers specific for mouse Gapdh, Cyp1a1, Cyp1a2, and Cyp1b1. Gapdh was used as an endogenous control. Primer sequences are as follows:

*Gapdh:* Forward: 5′-CATCACTGCCACCCAGAAGACTG-3′

Reverse: 5′-ATGCCAGTGAGCTTCCCGTTCAG-3′

*Cyp1a1:* Forward: 5′-CCTCATGTACCTGGTAACCA-3′

Reverse: 5′-AAGGATGAATGCCGGAAGGT-3′

*Cyp1a2:* Forward: 5′-CATCACAAGTGCCCTGTTCAAGC-3′

Reverse: 5′-AATGCTCCAGGTGATGGCTGTG-3′

*Cyp1b1:* Forward: 5′-GCCACTATTACGGACATCTTCGG-3′

Reverse: 5′-ACAACCTGGTCCAACTCAGCCT-3′

*Nppa* Forward: 5′-GAACCTGCTAGACCACCT-3′

Reverse: 5′-CCTAGTCCACTCTGGGCT-3′

*Nppb* Forward: 5′-AAGCTGCTGGAGCTGATAAGA-3′

Reverse: 5′-GTTACAGCCCAAACGACTGAC-3′

*Fundc1* Forward: 5′-CCCCCTCCCCAAGACTATGAA-3′

Reverse: 5′-CCACCCATTACAATCTGAGTAGC-3′

*Ccl21c* Forward: 5′-AAGGCAGTGATGGAGGGGGA-3′

Reverse: 5′-GGCTTAGAGTGCTTCCGGGGTA-3′

### Statistical analysis

Results are presented as mean ± SEM. Comparisons between two groups were conducted using unpaired two-tailed Student t-tests. Comparisons among four groups were conducted using one-way or two-way analysis of variance (ANOVA) followed by a post-hoc test, such as Tukey’s HSD or Bonferroni correction. A level of p ≤ 0.05 was considered to be significant, and “ns” (non-significant) denotes p > 0.05 between groups.

## RESULTS

### Cyp1 Enzymes Are Differentially Expressed Following Dox Treatment in Mice

To evaluate the involvement of Cyp1 enzymes in Dox-induced cardiomyopathy, we first examined the expression of *Cyp1a1*, *Cyp1a2*, and *Cyp1b1* by reverse transcription-quantitative polymerase chain reaction (RT-qPCR) in the liver and myocardium of mice treated with saline or Dox, using a low-dose Dox regimen that phenocopies the chronic toxicity observed in human patients.^39,41^ To assess the expression of *Cyp1* shortly after administering Dox, mice were sacrificed within 24 hours of the last dose of Dox, and hearts and livers were collected. All *Cyp1* genes were expressed in both tissues. *Cyp1a1* remained unchanged in the liver and heart for both male and female mice treated with saline or Dox (**Figure 1A and B**). *Cyp1a2* was downregulated in the liver for both male and female mice treated with Dox compared to those treated with saline (**Figure 1C**). In the heart, this effect was more consistently observed in female mice (**Figure 1D**). In contrast, *Cyp1b1* was markedly upregulated in both the liver and heart in response to Dox, particularly in male mice (**Figure 1E and F**). These data indicate that chronic, low-dose Dox treatment modulates expression of *Cyp1a2* and *Cyp1b1* in the liver and the heart.

**Figure 1.**
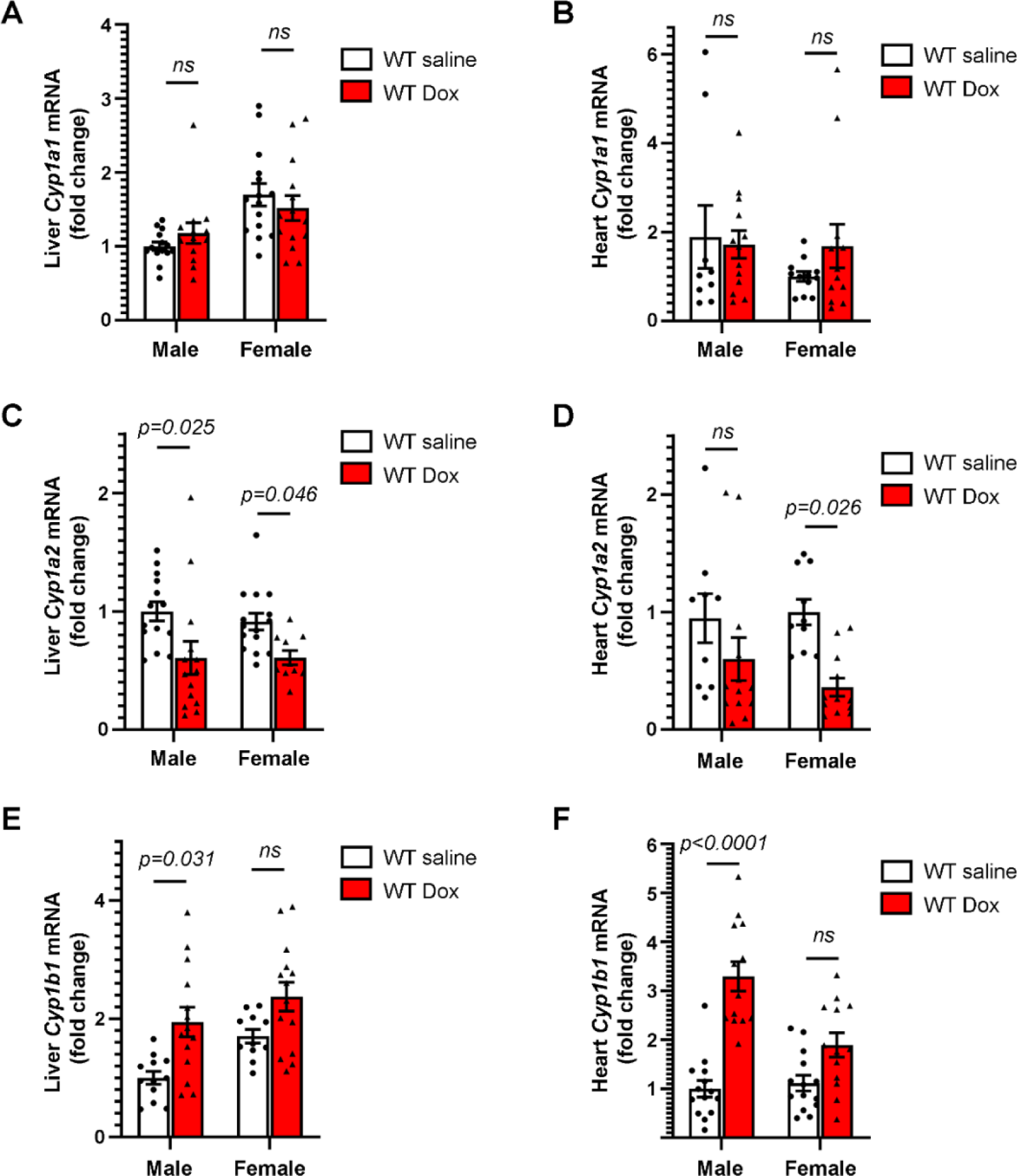
Cyp1 isoforms are differentially expressed in the context of Dox treatment. Mice aged 12-14 weeks were treated with saline or Dox every other day for 2 weeks, totaling 8 injections.^39^ Tissues were obtained within 24 hours after the last Dox treatment. Gene expression of *Cyp1a1* in the liver (**A**) or heart (**B**), *Cyp1a2* in the liver (**C**) or heart (**D**), and *Cyp1b1* in the liver (**E**) or heart (**F**), was determined by RT-qPCR in male or female mice. n = 12-15 mice/group. Results for all groups were normalized to GAPDH levels and expressed relative to the saline-treated control of the same gender. Values are presented as mean ± SEM. Unpaired two-tailed Student t-tests were used to compare the differences between the saline-treated group and the Dox-treated group within the same gender and tissue.

### Chemical Inhibition of Cyp1 Prevents Dox Cardiomyopathy in Mice

YW-130, a visnagin analog and a novel Cyp1 inhibitor synthesized by our group, was previously found to protect against acute Dox cardiotoxicity in zebrafish and mice at low doses with minimal toxicity.^35^ To guide our experimental design, we first measured the half-life of YW-130 in mouse plasma to be approximately 40 minutes when delivered by the i.p. route. To investigate the contribution of Cyp1 enzymes to the development of chronic Dox cardiomyopathy in mice, we treated WT mice with saline, YW-130 (1 mg/kg i.p. every other day for 2 weeks), Dox (3 mg/kg i.p. every other day for 2 weeks, totaling 24 mg/kg), and a combination of YW-130 and Dox, as summarized in **Figure 2A**. To evaluate the delayed cardiac dysfunction induced by a repeated low doses of Dox, transthoracic echocardiography was performed to assess cardiac function 7-10 weeks after the initiation of Dox treatment, followed by tissue harvesting. As expected, Dox treatment resulted in decreased cardiac function, as demonstrated by reduced fractional shortening (FS)% and increased left ventricular internal diameter at end-systole (LVIDs). Co-treatment with YW-130 prevented cardiac dysfunction in mice treated with Dox (**Figure 2B-E**). Notably, co-treatment with YW-130 did not affect the heart weight to tibia length ratio (HW/TL) or change in bodyweight (**Figure 2F and 2G**).

**Figure 2.**
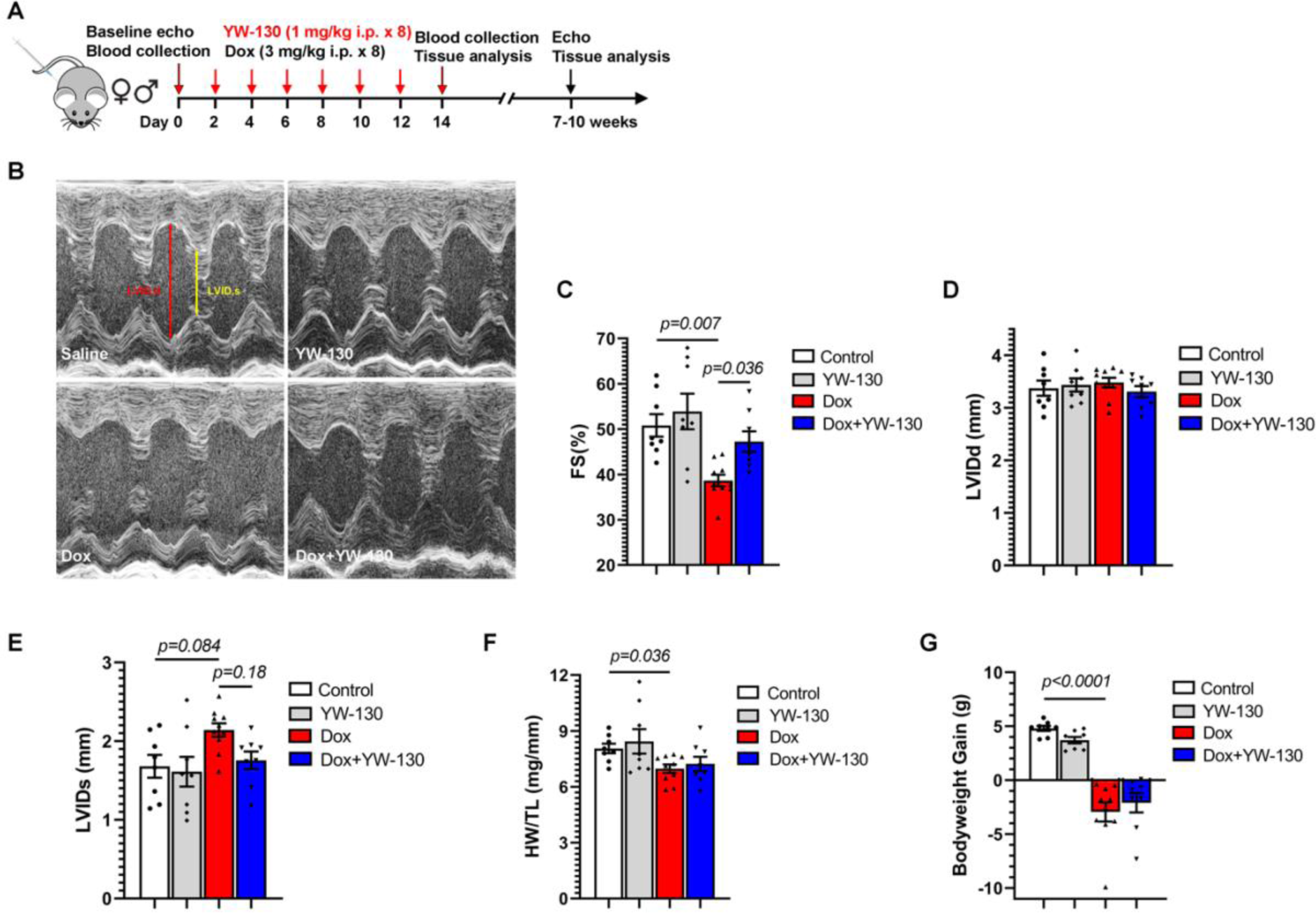
YW-130 treatment prevents Dox-induced cardiac dysfunction. (**A**) Summary of mouse treatment protocol. (**B**) Representative echocardiography images in mice treated with saline, YW-130, Dox, and the combination of YW-130 and Dox. Cardiac function parameters were assessed by echocardiography, including percentage fractional shortening (FS%) (**C**), left ventricular internal diameter at end-diastole (LVIDd) (**D**), and left ventricular internal diameter at end-systole (LVIDs) (**E**). (**F**) Ratio of heart weight (HW) to tibia length (TL). (**G**) Body weight gain from pre-treatment until pre-harvest. n=8, 8, 10, 8 in the groups receiving saline, YW-130, Dox, and the combination of YW-130 and Dox treatment, respectively. Values are presented as mean ± SEM. One-way ANOVA was used to compare the differences between groups, followed by Tukey’s HSD post-hoc test.

#### Knockout of Cyp1 Genes Protects Mice Against Dox Cardiomyopathy

To further evaluate the protective role of Cyp1 inhibition in Dox cardiomyopathy, we generated Cyp1 triple-knockout mice by breeding *Cyp1a1/1a2^-/-^* mice with *Cyp1b1^-/-^* mice for several generations. *Cyp1a1/1a2/1b1^-/-^* mice did not have altered lifespan or reproductive capacity. Following confirmation of strain identity and the establishment of stable lines, we treated WT or *Cyp1a1/1a2/1b1^-/-^* mice with either saline or Dox (5 mg/kg i.v.) once weekly for a total of 4 weeks, as illustrated in **Figure 3A**.^40^ Notably, *Cyp1a1/1a2/1b1^-/-^* mice exhibited higher FS% at baseline compared to WT mice (**Figure 3B**), suggesting a role for Cyp1 enzymes in normal cardiovascular homeostasis. Seven to ten weeks after the initiation of Dox treatment, *Cyp1a1/1a2/1b1^-/-^* mice were protected from chronic Dox-induced cardiac dysfunction, as assessed by echocardiography (**Figure 3C-G**). This effect was more pronounced in male mice, potentially due to lower baseline FS% in males compared to females as previously described.^42^

**Figure 3.**
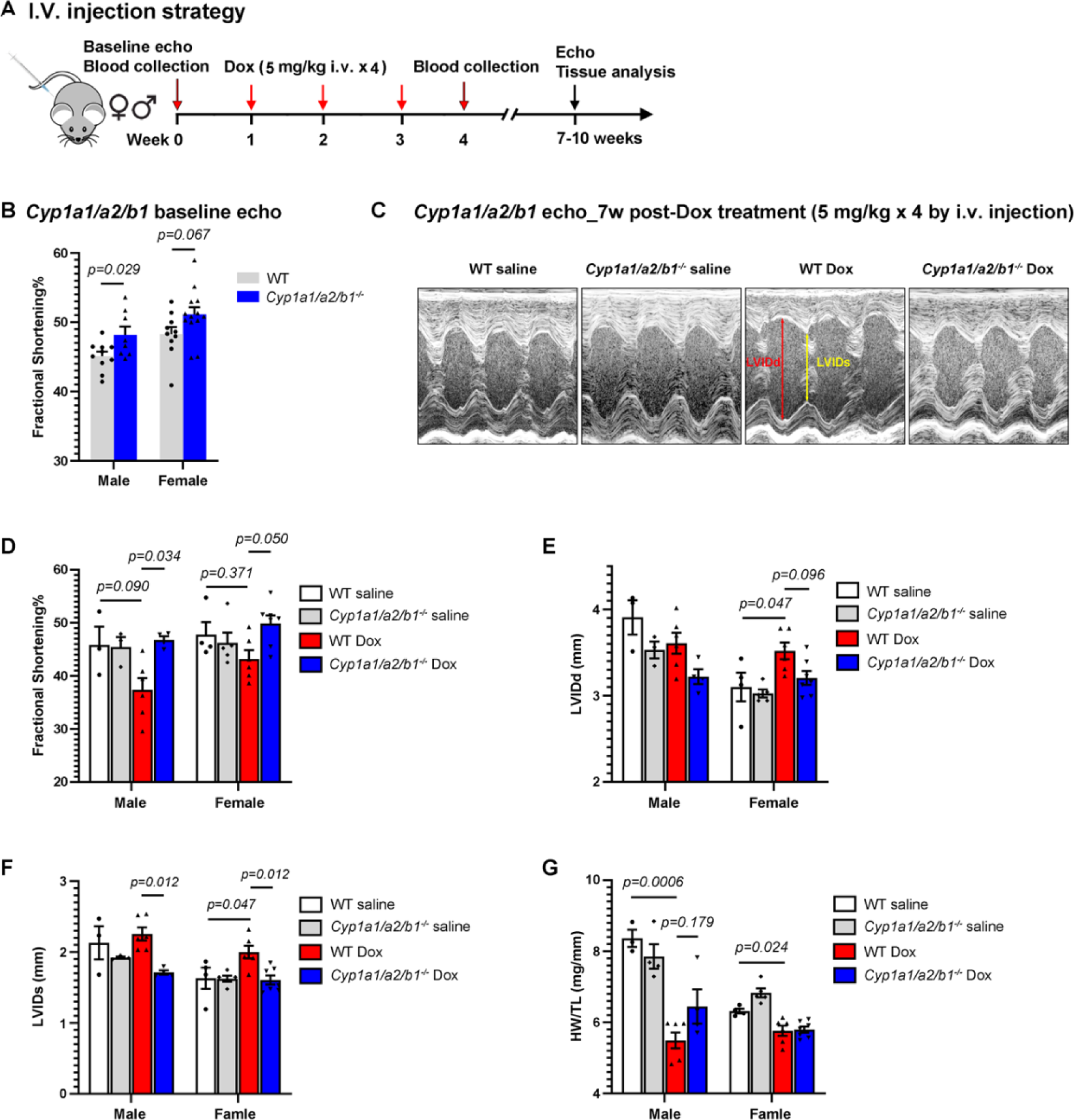
Knockout of *Cyp1a1/1a2/1b1* protects against Dox-induced cardiac dysfunction. (**A**) Summary of mouse treatment protocol. (**B**) Percent fractional shortening (FS%) of male and female WT or *Cyp1a1/1a2/1b1^-/-^* mice at baseline; n=8-13/group. (**C**) Representative echocardiography images in WT or *Cyp1a1/1a2/1b1^-/-^*mice treated with saline or Dox, measured 7 weeks after the initiation of Dox treatment. Cardiac function parameters were assessed by echocardiography, including percentage fractional shortening (FS%) (**D**), left ventricular internal diameter at end-diastole (LVIDd) (**E**), and left ventricular internal diameter at end-systole (LVIDs) (**F**). n=3-4 in saline-treated groups and n=3-7 in Dox-treated groups. (**G**) Ratio of heart weight (HW) to tibia length (TL). n=3-7/group. Values are presented as mean ± SEM. Two-way ANOVA was used to compare the differences between groups, followed by Bonferroni correction post-hoc test

### Knockout of Cyp1a Enzymes Confers Cardioprotection in Male and Female Mice, Whereas Knockout of Cyp1b1 Exacerbates Dox Cardiomyopathy in Male Mice

To explore how inhibition of specific Cyp1 isoforms contributes to cardioprotection in the context of Dox cardiomyopathy, we administered Dox to WT, *Cyp1a1/1a2^-/-^*, and *Cyp1b1^-/-^* mice a shown in **Figure 3A**. Male and female *Cyp1a1/1a2^-/-^* mice were protected from chronic Dox-induced cardiac dysfunction (**Figure 4**), as demonstrated by improved FS%. As expected, Dox treatment led to upregulation of cardiac *Nppa* and *Nppb*, molecule markers of cardiac injury and stress. This effect was mitigated in male *Cyp1a1/1a2^-/-^* mice (**Supplementary Figure 1**). Conversely, male *Cyp1b1^-/-^* mice exhibited increased cardiac dysfunction following Dox treatment (**Figure 5**) compared to WT mice treated with Dox. In female mice, deletion of *Cyp1b1* had no significant effect on cardiac function (**Figure 5**). As in triple-knockout mice, similar changes in baseline cardiac function were observed in the absence of treatment.

**Figure 4.**
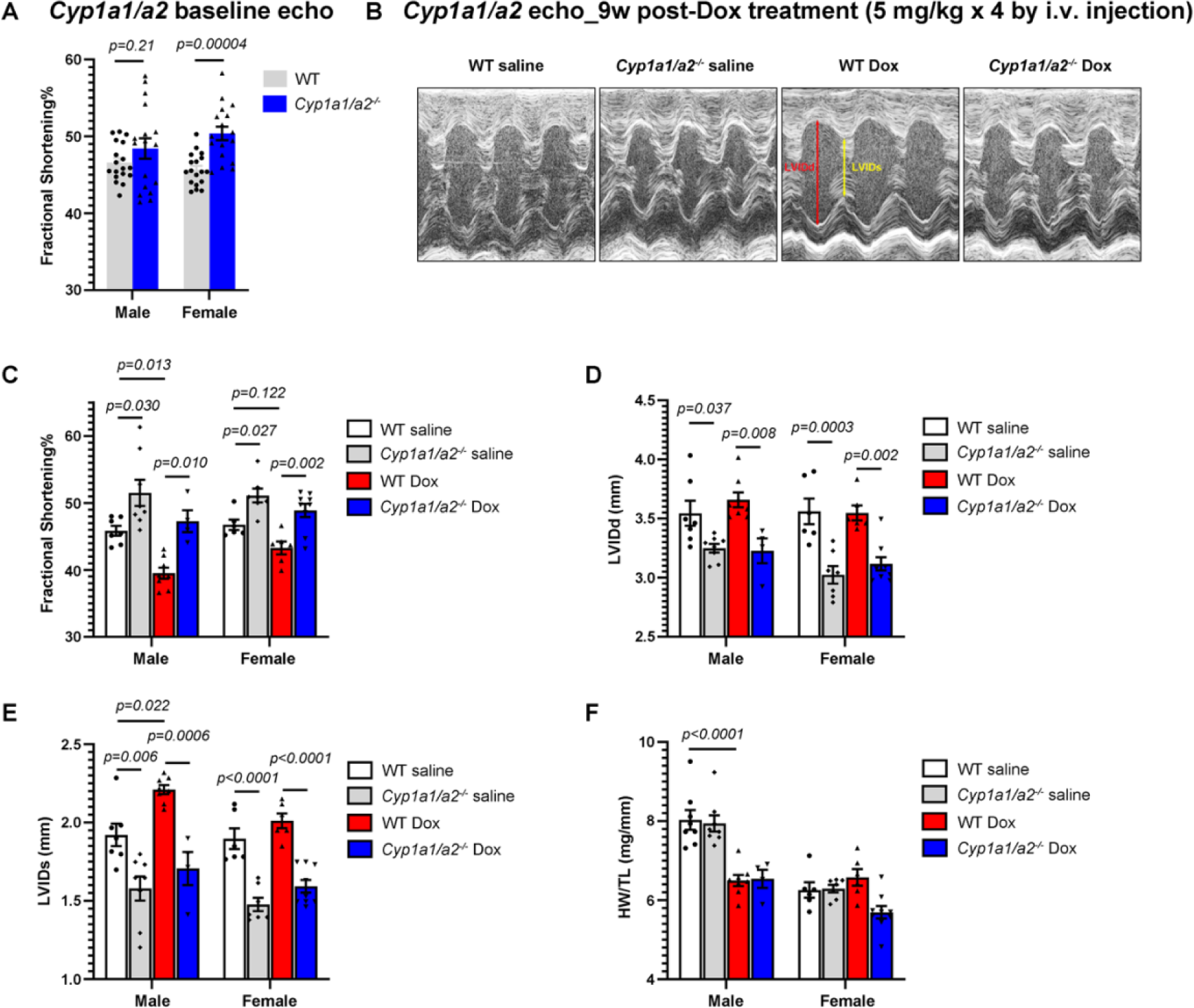
*Cyp1a* deficiency protects against Dox-induced cardiac dysfunction. (**A**) Percent fractional shortening (FS%) of male and female WT or *Cyp1a1/1a2^-/-^* mice at baseline; n=16-18/group. (**B**) Representative echocardiography images in WT or *Cyp1a1/1a2^-/-^* mice treated with saline or Dox, measured 7 weeks after the initiation of Dox treatment. Cardiac function parameters were assessed by echocardiography, including percentage fractional shortening (FS%) (**C**), left ventricular internal diameter at end-diastole (LVIDd) (**D**), and left ventricular internal diameter at end-systole (LVIDs) (**E**). n=4-9 mice/group. (**F**) Ratio of heart weight (HW) to tibia length (TL). n=4-8 mice/group. Values are presented as mean ± SEM. Two-way ANOVA was used to compare the differences between groups, followed by Bonferroni correction post-hoc test.

**Figure 5.**
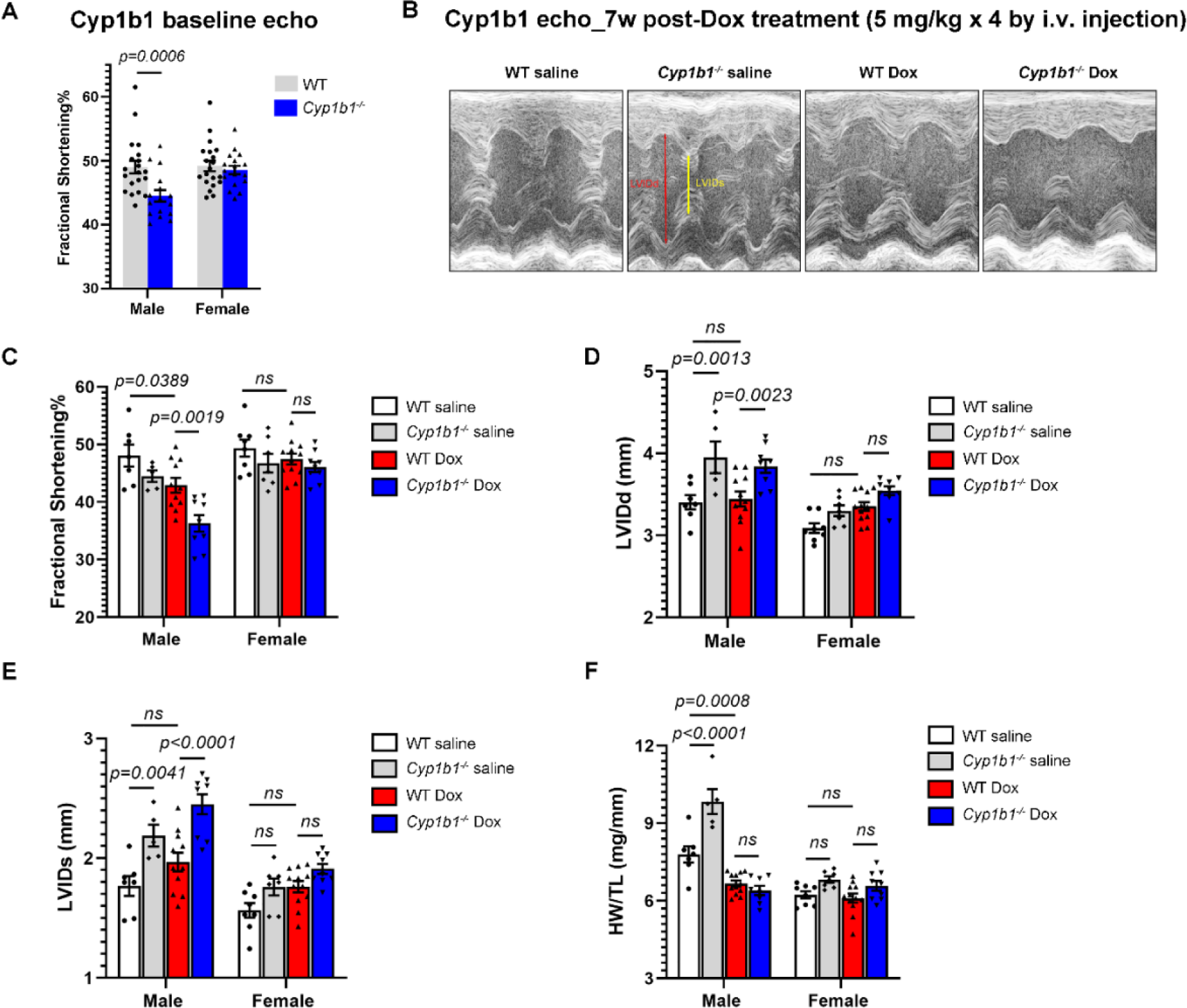
*Cyp1b1* deficiency exacerbates Dox-induced cardiac dysfunction in male mice. (**A**) Percent fractional shortening (FS%) of male and female WT or *Cyp1b1^-/-^* mice at baseline; n=15-20/group. (**B**) Representative echocardiography images in male WT or *Cyp1b1^-/-^* mice treated with saline or Dox, measured 7 weeks after the initiation of Dox treatment. Cardiac function parameters were assessed by echocardiography, including percentage fractional shortening (FS%) (**C**), left ventricular internal diameter at end-diastole (LVIDd) (**D**), and left ventricular internal diameter at end-systole (LVIDs) (**E**). n=5-11 mice/group. (**F**) Ratio of heart weight (HW) to tibia length (TL). n=5-11 mice/group. Values are presented as mean ± SEM. Two-way ANOVA was used to compare the differences between groups, followed by Tukey’s HSD post-hoc test.

### RNA Sequencing Reveals Fundc1 and Ccl21c as Potential Molecular Targets in Cardioprotection Mediated by Cyp1a

To explore the mechanisms underlying the protective effects of Cyp1a inhibition in Dox cardiomyopathy, we administered saline or Dox to both WT mice and *Cyp1a1/1a2^-/-^* mice using the previously described Dox regimen (Dox 3 mg/kg, i.p., every other day for two weeks; **Figure 2A**). To investigate the early molecular changes that precede cardiac dysfunction, heart tissues were collected one day after the last injection, and the total RNA extracted from left ventricular myocardial tissue was subjected to bulk RNAseq analysis (**Supplementary Data**). As expected, we observed significant differences in transcriptomes between mice treated with saline and those treated with Dox. A total of 3,233 genes were found to be differentially expressed in Dox-treated mice, with 1,864 genes showing increased expression and 1,369 genes showing decreased expression (adjusted p < 0.05). The top 500 differentially expressed genes showed enrichment in pathways involved in mitochondrial function, cellular respiration, and oxidative stress associated with Dox treatment (**Supplementary Figure 2**). These results are consistent with previous reports of altered energy metabolism and oxidative stress as key contributors to Dox-induced cardiotoxicity.

Comparing *Cyp1a*-deficient mice to WT mice, both treated with Dox, we identified changes in autophagy and mitophagy associated with *Cyp1a* knockout through biological and reactome analyses (**Figure 6A-B**). Mitophagy is a specialized form of autophagy responsible for selectively removing damaged or dysfunctional mitochondria, particularly in response to metabolic stress conditions, contributing to mitochondrial homeostasis.^43^ Specifically in this pathway, we observed upregulation of the gene FUN14 domain-containing 1 (*Fundc1*) in *Cyp1a1/1a2^-/-^* mice compared to WT mice. These findings were observed with saline treatment (log fold change (FC)=0.92, adjusted p=9.25E-5) and with Dox treatment (log FC=2.08, adjusted p=0.003), as validated by RT-qPCR analysis (**Figure 6C**). These findings were not observed when comparing *Cyp1a1/1a2/1b1^-/-^* mice to WT mice.

**Figure 6.**
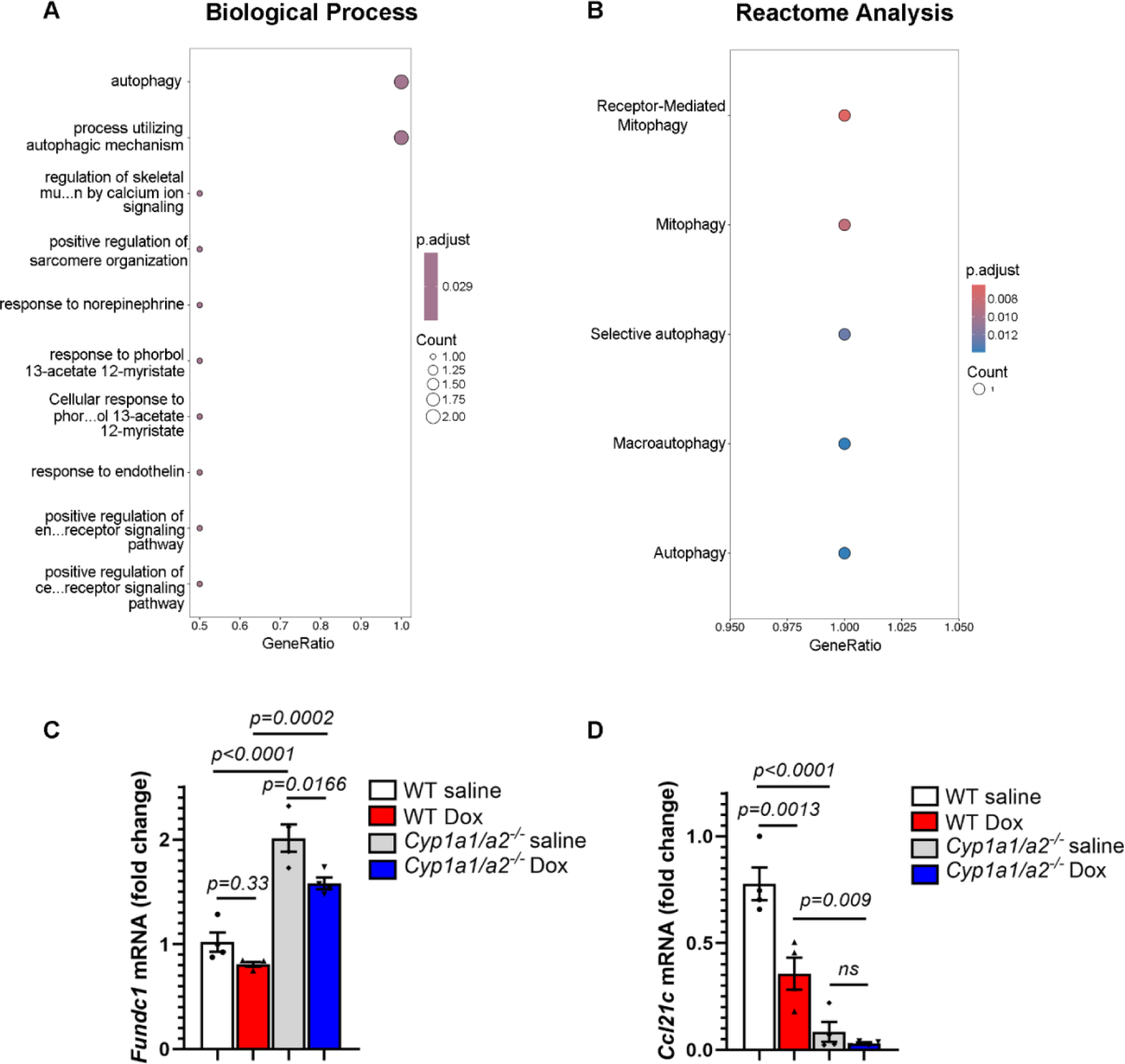
RNA Sequencing Reveals *Fundc1* and *Ccl21c* as Potential Mediators of Cardioprotection Conferred by *Cyp1a* Inhibition. (**A**) Biological pathway analysis of *Cyp1a1/1a2^-/-^* mice treated with Dox compare to WT mice treated with Dox. (**D**) Reactome analysis of *Cyp1a1/1a2^-/-^* mice treated with Dox compare to WT mice treated with Dox. n=3-4 mice/group. mRNA levels of (**E**) *Fundc1* and (**F**) *Ccl21c* in mouse hearts of WT or *Cyp1a1/1a2^-/-^* mice treated with saline or Dox, respectively, as measured by RT-qPCR. Values are presented as mean ± SEM. One-way ANOVA was used to compare the differences between groups, followed by Tukey’s HSD post-hoc test.

We also observed downregulation of C-C motif chemokine 21c (*Ccl21c),* a chemokine involved in immune cell trafficking and activation,^44,45^ in *Cyp1a1/1a2^-/-^* mice treated with either saline or Dox. This finding was also validated by RT-qPCR analysis (**Figure 6D**). Similar downregulation of *Ccl21c* was also observed in *Cyp1a1/1a2/1b1^-/-^* mice with saline or Dox treatment, and in WT mice treated with Dox and YW-130 (**Supplementary Data**), suggesting that *Ccl21c* may contribute to the cardioprotective effects of *Cyp1a* inhibition.

## DISCUSSION

Anthracycline-induced cardiomyopathy and heart failure confer a poor prognosis in patients with cancer.^2^ This study explored the potential therapeutic role of inhibiting Cyp1 in the context of a mouse model that recapitulates the chronic cardiomyopathy seen in patients. Our findings provide insights into the differential functions of Cyp1 isoforms in Dox-induced cardiac dysfunction. Specifically, we found that *Cyp1a1/1a2^-/-^* and *Cyp1a1/1a2/1b1^-/-^* mice were protected from Dox cardiomyopathy, similar to mice treated with the CYP inhibitor YW-130. In contrast, *Cyp1b1^-/-^* mice demonstrated worse cardiac function after Dox treatment. Taken together, these data support inhibition of *Cyp1a* as a promising cardioprotective strategy in patients treated with Dox.

This study assessed cardiac *Cyp1* expression levels in the heart at an early time point, within 24 hours after the last Dox treatment. We observed reduced *Cyp1a2* expression without significant changes in *Cyp1a1* levels. These data differ from a prior report from our group, where a decrease in *Cyp1a1* expression was observed in female C57BL/6 mice 24 hours after treatment with a single high dose of Dox (20 mg/kg i.p.).^29^ Moreover, Dox caused induction of both *Cyp1a1* and *Cyp1a2* in H9c2 cells in vitro.^46^ Several factors could contribute to the discordance between these studies, including the timing of gene expression assessment and the use of a single high-dose Dox regimen.

On the other hand, Dox-mediated induction of *Cyp1b1* has been consistently observed in both *in vitro* and *in vivo* studies, in zebrafish and in mice.^29,35,46^ Acute and chronic administration of Dox has been shown to upregulate *Cyp1b1* expression in various organs, including the heart, liver, and kidney.^29,47,48^ Our observations align with other reports suggesting a sex-specific response, where acute Dox administration (20 mg/kg i.p.) was associated with the induction of cardiac *Cyp1b1* in male but not female C57Bl/6 mice^29^. Male mice also demonstrated more pronounced cardiac dysfunction in this model.

Dox has been shown to activate the AhR and therefore AhR-regulated genes such as *Cyp1a1* and *Cyp1b1*, in both H9c2 cells and in the rat heart.^30,34^ However, *Cyp1a1* is also regulated by the AhR but was not induced in our study and others,^49^ suggesting that other regulatory mechanisms beyond the AhR pathway could contribute to *Cyp1* induction. One potential factor could be cardiac inflammation, which has been shown to inhibit *Cyp1a1* and *Cyp1a2* while inducing *Cyp1b1.*^50,51^ For instance, previous studies have reported a significant induction of *Cyp1b1* in the liver of male rats in response to lipopolysaccharide-induced inflammation, an effect that was attributed to androgen signaling.^51^ In prior work, induction of the AhR was shown to be protective in rats treated with Dox,^34^ an effect that may be related to the pleiotropic nature of the AhR in regulating expression of both protective and deleterious pathways, in addition to its described function as a transcriptional regulator of *Cyp1*. Our data suggest that inhibition of downstream *Cyp1* enzymes may serve as a more nuanced cardioprotective strategy, compared to inhibition of the AhR itself.

Our RNAseq analyses highlight two potential molecular targets of *Cyp1a*-mediated cardioprotection. *Fundc1* is a mitophagy receptor localized to mitochondria-associated membranes (MAMs).^52,53^ It interacts with other proteins and cellular structures, such as LC3 proteins and the endoplasmic reticulum, to promote the engulfment of damaged mitochondria by autophagosomes, facilitating their removal from the cell.^54^ This process is essential for maintaining cellular homeostasis and preventing the accumulation of dysfunctional mitochondria, which can lead to cellular stress and dysfunction. Fundc1-triggered mitophagy has been shown to offer protection against ischemia-reperfusion (I/R) injury and Dox-induced cardiac dysfunction.^55–57^ Our data suggest that Fundc1-mediated mitophagy may be a potential mechanism through which *Cyp1a* inhibition mitigates Dox-induced cardiomyopathy. Wu et al. previously reported that deficiency of *Fundc1* in mice impairs heart function.^54^ *Fundc1* deficiency aggravated Dox-induced cardiac dysfunction, mitochondrial injury, and cardiomyocyte PANoptosis (pyroptosis, apoptosis, necrosis) in mice,^57^ suggesting a cardioprotective role of *Fundc1* in Dox cardiomyopathy. Furthermore, this study reported downregulation of *Fundc1* in myocardial tissue from patients with dilated cardiomyopathy, suggesting that these findings could potentially be translated to the clinic.

Inflammation has been implicated in Dox-induced cardiotoxicity.^58–60^ C-C motif chemokine ligand 21c (Ccl21c), a chemokine involved in immune cell trafficking, has been implicated in atherosclerosis, myocardial infarction, and heart failure in patients.^61–63^ Increased levels of CCL21 have been detected in ruptured lesions in the coronary arteries of patients with myocardial infarction^64^ and in explanted myocardial tissue from patients with heart failure.^62^ Interestingly, although Dox treatment has been associated with a robust inflammatory response in the heart, WT mice treated with Dox demonstrated downregulation of Ccl21c. Additional studies will be necessary to define the mechanisms regulating expression of this chemokine in both WT and *Cyp1a*-deficient mice.

### Limitations

This study leverages mouse models to provide valuable insights into the role of Cyp1 isoforms in anthracycline-induced cardiomyopathy. Additional mechanistic studies are ongoing to elucidate the role of Fundc1-mediated mitophagy, Ccl21c-mediated inflammation, and other molecular pathways that could contribute to Cyp1-mediated cardioprotection. Furthermore, our current work utilized mouse models with ubiquitous deletion of *Cyp1*; development of mouse strains lacking *Cyp1a1* or *Cyp1a2* in specific cell types (e.g., cardiomyocytes or hepatocytes) is ongoing. Finally, while inhibition of *Cyp1a* was cardioprotective in our experiments, further investigation will be necessary to elucidate the mechanistic role of *Cyp1b1* in Dox cardiomyopathy.

## CONCLUSION

This study highlights the critical role of *Cyp1* isoforms in Dox-induced cardiomyopathy in mice. Both pharmacologic inhibition of *Cyp1* through YW-130 treatment and genetic deficiency of *Cyp1a1/1a2* effectively prevented Dox-induced cardiac toxicity. These findings highlight the potential therapeutic potential of *Cyp1a* inhibition in preventing anthracycline-induced cardiac toxicity. Additionally, the study provides valuable insights into potential mechanisms by which *Cyp1* inhibition confers cardioprotection.

## Acknowledgments

The authors thank Dr. Daniel W. Nebert, MD, from the University of Cincinnati College of Medicine, and Dr. Frank J. Gonzalez, Ph.D, from the National Cancer Institute, for kindly providing the *Cyp1a1/1a2^-/-^* and *Cyp1b1^-/-^* mouse founders, respectively.

## Nonstandard Abbreviations and Acronyms

Dox: Doxorubicin
Cyp1: Cytochrome P450 family 1
RT-qPCR: Reverse transcription-quantitative polymerase chain reaction
WT: Wild-type
i.p.: Intraperitoneal
i.v.: Intravenous
NRF2: Nuclear factor erythroid 2-related factor 2
AMPK: AMP-activated protein kinase
PI3K/Akt: Phosphatidylinositol 3-kinase/protein kinase B
GO: Gene ontology
EF: Ejection fraction
FS: Fractional shortening
LVIDd: Left ventricular internal diameter at end-diastole
LVIDs: Left ventricular internal diameter at end-systole
HW/TL: Heart weight / tibia length
ANOVA: Analysis of variance
AhR: Aryl hydrocarbon receptor
Nppa: Natriuretic peptide A
Nppb: Natriuretic peptide B
Fundc1: FUN14 domain-containing protein 1
Ccl21: C-C motif chemokine ligand 21c

