## Supplementary Figures for "Inhibition of Cyp1a Protects Mice against Anthracycline Cardiomyopathy"

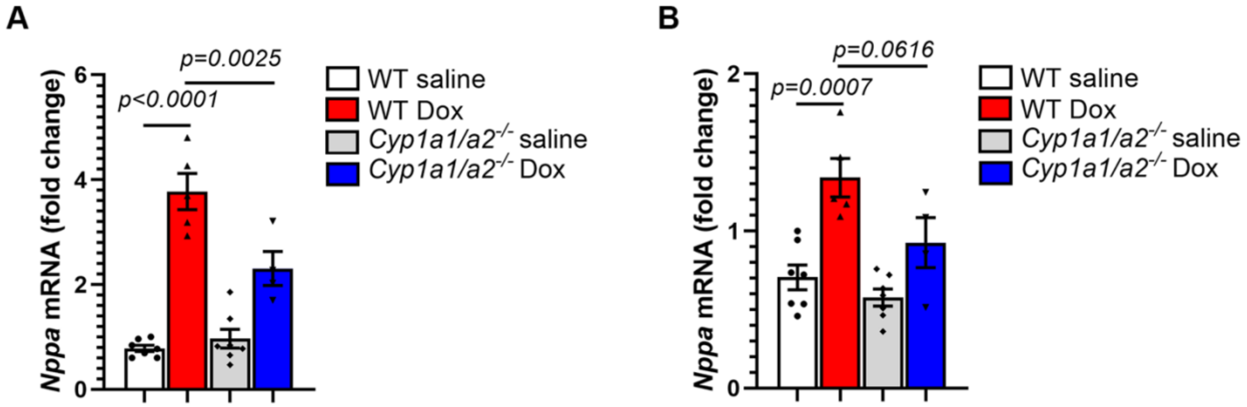


**Supplementary Figure 1. *Cyp1a*-null mice have decreased expression of cardiac natriuretic peptides following treatment with Dox.** mRNA levels of (**A**) *Nppa* and (**B**) *Nppb* in mouse hearts of WT or *Cyp1a1/1a2*$^{-/-}$ mice treated with saline or Dox, respectively, as measured by RT-qPCR. Values are presented as mean ± SEM. One-way ANOVA was used to compare the differences between groups, followed by Tukey's HSD post-hoc test.

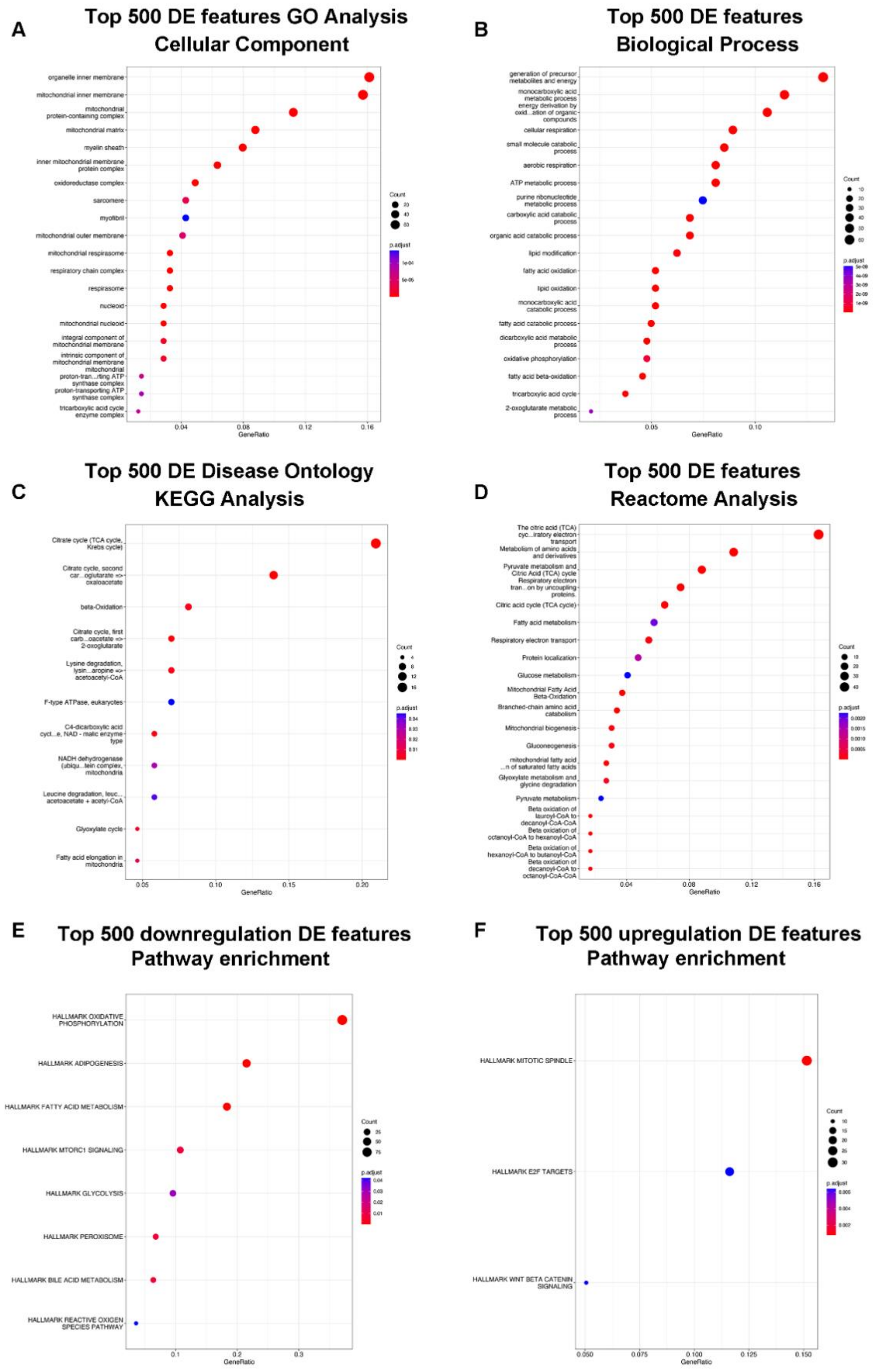
A
Top 500 DE features GO Analysis
Cellular Component
organelle inner membrane
mitochondrial inner membrane
mitochondrial protein-containing complex
mitochondrial matrix
myelin sheath
inner mitochondrial membrane protein complex
oxidoreductase complex
sarcomere
myofibril
mitochondrial outer membrane
mitochondrial respirasome
respiratory chain complex
respirasome
nucleoid
mitochondrial nucleoid
integral component of mitochondrial membrane
intrinsic component of mitochondrial membrane
mitochondrial proton-tran...rting ATP synthase complex
proton-transporting ATP synthase complex
tricarboxylic acid cycle enzyme complex
0.04
0.08
0.12
0.16
GeneRatio
Count
20
40
60
p.adjust
1e-04
5e-05
B
Top 500 DE features
Biological Process
generation of precursor metabolites and energy
monocarboxylic acid metabolic process
energy derivation by oxid...ation of organic compounds
cellular respiration
small molecule catabolic process
aerobic respiration
ATP metabolic process
purine ribonucleotide metabolic process
carboxylic acid catabolic process
organic acid catabolic process
lipid modification
fatty acid oxidation
lipid oxidation
monocarboxylic acid catabolic process
fatty acid catabolic process
dicarboxylic acid metabolic process
oxidative phosphorylation
fatty acid beta-oxidation
tricarboxylic acid cycle
2-oxoglutarate metabolic process
0.05
0.10
GeneRatio
Count
10
20
30
40
50
60
p.adjust
5e-09
4e-09
3e-09
2e-09
1e-09
C
Top 500 DE Disease Ontology
KEGG Analysis
Citrate cycle (TCA cycle, Krebs cycle)
Citrate cycle, second car...oglutarate => oxaloacetate
beta-Oxidation
Citrate cycle, first carb...oacetate => 2-oxoglutarate
Lysine degradation, lysin...aropine => acetoacetyl-CoA
F-type ATPase, eukaryotes
C4-dicarboxylic acid cycl...e, NAD - malic enzyme type
NADH dehydrogenase (ubiqu...tein complex, mitochondria
Leucine degradation, leuc... acetoacetate + acetyl-CoA
Glyoxylate cycle
Fatty acid elongation in mitochondria
0.05
0.10
0.15
0.20
GeneRatio
Count
4
8
12
16
p.adjust
0.04
0.03
0.02
0.01
D
Top 500 DE features
Reactome Analysis
The citric acid (TCA) cyc...iratory electron transport
Metabolism of amino acids and derivatives
Pyruvate metabolism and Citric Acid (TCA) cycle
Respiratory electron tran...on by uncoupling proteins.
Citric acid cycle (TCA cycle)
Fatty acid metabolism
Respiratory electron transport
Protein localization
Glucose metabolism
Mitochondrial Fatty Acid Beta-Oxidation
Branched-chain amino acid catabolism
Mitochondrial biogenesis
Gluconeogenesis
mitochondrial fatty acid ...n of saturated fatty acids
Glyoxylate metabolism and glycine degradation
Pyruvate metabolism
Beta oxidation of lauroyl-CoA to decanoyl-CoA-CoA
Beta oxidation of octanoyl-CoA to hexanoyl-CoA
Beta oxidation of hexanoyl-CoA to butanoyl-CoA
Beta oxidation of decanoyl-CoA to octanoyl-CoA-CoA
0.04
0.08
0.12
0.16
GeneRatio
Count
10
20
30
40
p.adjust
0.0020
0.0015
0.0010
0.0005
E
Top 500 downregulation DE features
Pathway enrichment
HALLMARK OXIDATIVE PHOSPHORYLATION
HALLMARK ADIPOGENESIS
HALLMARK FATTY ACID METABOLISM
HALLMARK MTORC1 SIGNALING
HALLMARK GLYCOLYSIS
HALLMARK PEROXISOME
HALLMARK BILE ACID METABOLISM
HALLMARK REACTIVE OXIGEN SPECIES PATHWAY
0.1
0.2
0.3
GeneRatio
Count
25
50
75
p.adjust
0.04
0.03
0.02
0.01
F
Top 500 upregulation DE features
Pathway enrichment
HALLMARK MITOTIC SPINDLE
HALLMARK E2F TARGETS
HALLMARK WNT BETA CATENIN SIGNALING
0.050
0.075
0.100
0.125
0.150
GeneRatio
Count
10
15
20
25
30
p.adjust
0.006
0.004
0.002

**Supplementary Figure 2. RNA sequencing analysis of WT mice treated with saline or Dox.** (**A**) Cellular component analysis of the top 500 differentially expressed genes in mice treated with saline or Dox. (**B**) Biological process analysis of the top 500 differentially expressed genes in mice treated with saline or Dox. (**C**) KEGG analysis of the top 500 differentially expressed genes in mice treated with saline or Dox. (**D**) Reactome analysis of the top 500 differentially expressed genes in mice treated with saline or Dox. (**E**) Pathway analysis of top 500 downregulated differentially expressed genes in mice treated with saline or Dox. (**F**) Pathway analysis of top 500 upregulated differentially expressed genes in mice treated with saline or Dox. Values are presented as mean ± SEM. One-way ANOVA was used to compare the differences between groups, followed by Tukey’s HSD post-hoc test.
